# Subtype-Specific Roles of Nigrostriatal Dopaminergic Neurons in Motor and Associative Learning

**DOI:** 10.1101/2025.07.03.663035

**Authors:** Ahsan Habib, Gavin Riccobono, Lulu Tian, Disa Basu, Lixin Sun, Lisa Chang, Victor M. Martinez Smith, Lupeng Wang, Weidong Le, Huaibin Cai

## Abstract

Nigrostriatal dopaminergic neurons (DANs) in the *substantia nigra pars compacta* (SNc) comprise distinct subtypes defined by unique gene expression profiles and anatomical characteristics. However, their specific contributions to motor and non-motor functions remain elusive. Using *Calbindin 1* (*Calb1*) and *Aldehyde dehydrogenase* 1a1 (*Aldh1a1*) as molecular markers, we investigated the functions of these nigrostriatal DAN subtypes in mice. Through intersectional genetics and chemogenetic manipulation, we selectively inhibited *Calb1*^+^ or *Aldh1a1*^+^ DANs by stereotactically delivering an adeno-associated viral vector (AAV-CreOn-FlpOn-hM4Di-P2A-mCherry) into the SNc of *Th*^Flp^; *Calb1*^IRESCre^ or *Th*^Flp^; *Aldh1a1*^CreERT2^ double knock-in (KI) mice. This approach enabled subtype-specific neuronal inhibition via designer receptors exclusively activated by designer drugs (DREADD). Following DREADD ligand administration, both *Calb1^+^ and Aldh1a1^+^* DAN-inhibited mice exhibited significant reduction in voluntary movement and impaired motor skill learning, demonstrating their essential roles in motor function. However, only *Calb1*^+^ DAN inhibition affected early associative-learning behavior, suggesting a unique role in reinforcement learning. These findings establish *Calb1*^+^ and *Aldh1a1*^+^ nigrostriatal DANs as key regulators of movement and motor learning, with *Calb1*^+^ neurons additionally modulating reward-based associative learning. This study advances our understanding of the functional heterogeneity of nigrostriatal DAN subtypes and identifies potential therapeutic targets for addressing motor and non-motor deficits in Parkinson’s disease.

## Introduction

Parkinson’s disease (PD) is pathologically characterized by the widespread loss of midbrain dopaminergic neurons (DANs), especially those in the *substantia nigra pars compacta* (SNc) ^1–4^. We and others have demonstrated that Aldehyde dehydrogenase 1A1-positive (ALDH1A1^+^) DANs in the ventral tier of SNc are particularly vulnerable in individuals with PD, as well as in PD-related rodent and primate models ^5–9^. Recent single-cell and single-nuclei RNA sequencing studies have identified multiple molecularly distinct midbrain DAN subtypes ^10–12^. While midbrain DANs regulate various behavioral processes, including both motor and non-motor functions ^10,13–15^, the roles of specific DAN subtypes in regulating behaviors remain to be fully explored.

ALDH1A1^+^ DANs comprise two-thirds of the DANs in the human and rodent SNc ^5,16^ and one-third in the rodent ventral tegmental area (VTA) ^16^. In the SNc, these neurons contribute to locomotion control and are crucial for motor skill learning in rodents ^16^. While ALDH1A1^−^ DANs, located in the dorsal tier of the SNc, are also affected by PD, they are less vulnerable than ALDH1A1^+^ DANs ^5^. However, the role of ALDH1A1^−^ SNc DANs in motor control and learning remains unclear ^11,16,17^.

Since Calbindin 1 (CALB1) is predominantly expressed by ALDH1A1^−^ DANs ^9,17^, we employed an intersectional genetic approach using *Th*^Flp^ ^18^; *Calb1*^IRESTCre^ ^19^ double knock-in (KI) mice, combined with a Cre and Flp-dependent designer receptors exclusively activated by designer drugs (DREADD) system ^20^, to selectively inhibit *Calb1*^+^ DAN subtypes during behavioral tests. In parallel, we generated *Th*^Flp^; *Aldh1a1*^CreERT2^ ^16^ double KI mice to selectively manipulate the activity of *Aldh1a1*^+^ SNc DNAs. Our findings reveal a significant involvement of both *Aldh1a1*^+^ and *Calb1*^+^ SNc DANs in motor control and learning, highlighting both molecularly distinct SNc DAN subtypes as potential therapeutic targets for treating PD-related movement disorders.

## Results

### Selectively targeting of mouse *Calb1*^+^ SNc DANs using intersectional genetics and chemogenetic approaches

A recent study reported that the activity of *Calb1*^+^ SNc DANs is negatively correlated with movement acceleration ^17^. However, the direct impact of *Calb1*^+^ DANs on locomotion remains unclear. To investigate the function of this DAN subtype, we generated *Th*^Flp^; *Calb*1^IRESTCre^ (TC) double KI mice, enabling selective heterologous gene expression in *Calb1*^+^ DANs.

To specifically inhibit *Calb1*^+^ SNc DAN activity, we performed bilateral stereotactic injections of adeno-associated viral (AAV) vectors expressing either a Cre- and Flp-dependent inhibitory DREADD, hM4Di (AAV9-CreOn/FlpOn-hM4Di-P2A-mCherry), or a control red fluorescent protein mCherry (AAV9-CreOn/FlpOn-mCherry), into the SNc of TC double KI mice **(Fig. 1A)**. Immunostaining confirmed selective expression of hM4Di and mCherry in *Calb1*^+^ SNc DANs **(Fig. 1B)**, which are primarily located in the dorsal tier of SNc and account for approximately 30% of SNc DANs ^16^. Notably, when counterstaining sections with ALDH1A1 antibodies, we observed that while most mCherry-positive DANs were ALDH1A1-negative, a small fraction (5.8%) of mCherry-positive neurons in the dorsomedial SNc were also ALDH1A1-positive (arrows, **Fig. 1B, C**). Further examination of the projection pattern of *Calb1*^+^ SNc DANs in the dorsal striatum revealed that their axon terminals are primarily distributed in the dorsomedial striatum (DMS) **(Fig. 1D)**.

**Fig. 1.**
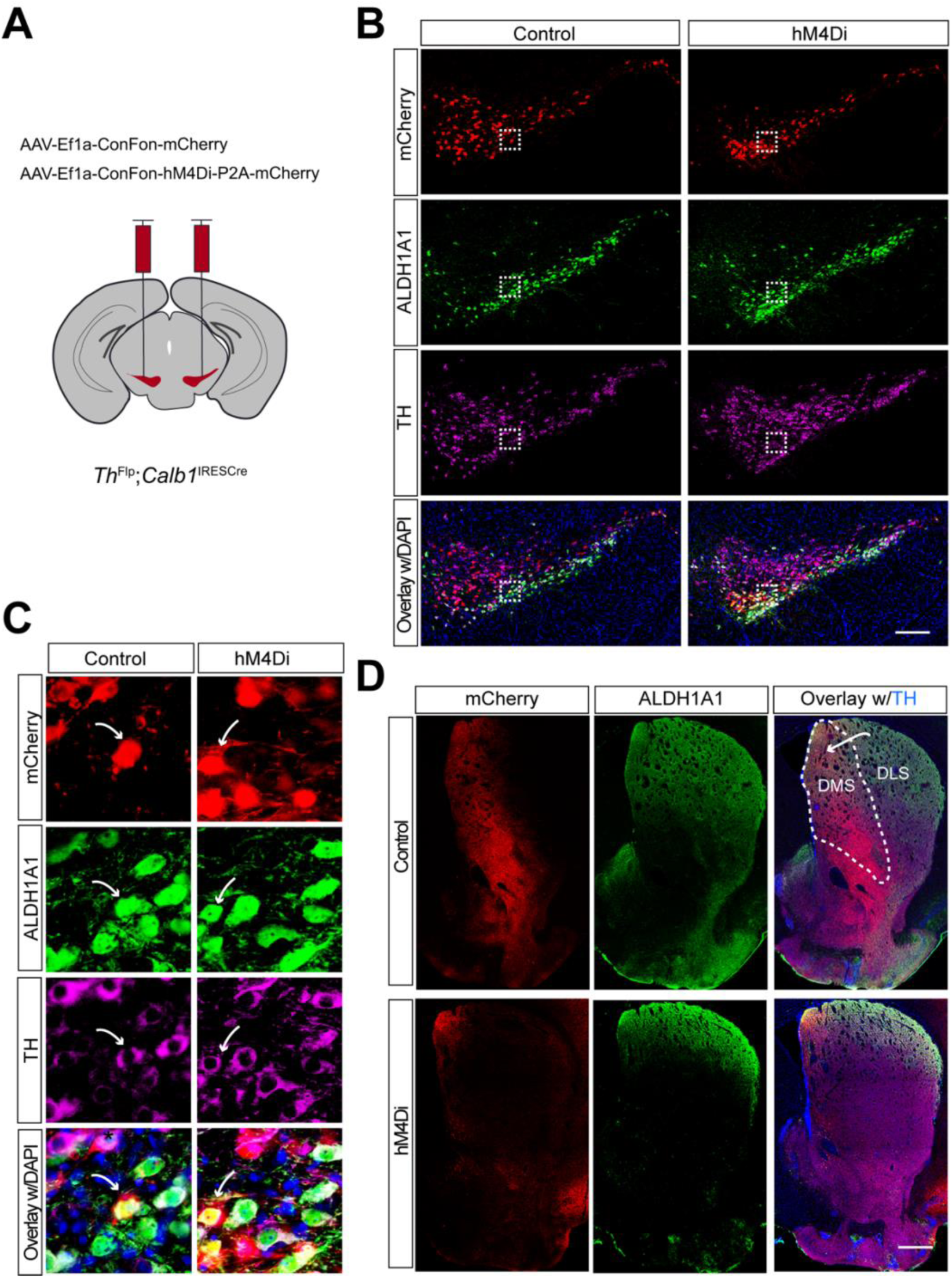
Targeted expression of the chemogenetic inhibitor hM4Di in *Calb1*^+^ nigrostriatal DANs. **(A)** Schematic depicting the stereotactic delivery of control mCherry or hM4Di AAVs in the SNc of *Th*^Flp^; *Calb1*^IRESCre^ (TC) double KI mice. (**B**) Representative images showing mCherry (red), ALDH1A1 (green), and TH (magenta) expression in the SNc of control mCherry (left panel) and hM4Di (right panel)-expressing mice. Scale bar: 1mm. (**C**) Magnified images of the boxed areas in (**B**), with arrows indicating neurons co-expressing mCherry and ALDH1A1. Scale bar: 100 μm. (**D**) Representative images illustrating mCherry (red), ALDH1A1 (green) and TH (magenta) staining in the striatum of control mCherry-expressing mice. Scale bar: 1mm.

Together, the use of TC double KI mice combined with local injection provides a precise approach to selectively targeting *Calb1*^+^ SNc DANs for genetic and functional manipulations.

### Chemogenetic inhibition of *Calb1*^+^ SNc DANs suppresses spontaneous movements

Six to eight weeks after stereotactic injection of control and DREADD AAVs, we used the beam break-based open-field test to assess spontaneous movements following intraperitoneal injection of the DREADD activator JHU37160 ^21^. We observed a marked reduction in total movement, ambulatory movement, rearing, and fine movement **(Fig. 2A-D)**. By contrast, intraperitoneal saline injections had no effect on locomotion or rearing **(Fig. 2E-H)**. These findings underscore the critical role of *Calb1*^+^ SNc DANs in regulating spontaneous movements.

**Fig. 2.**
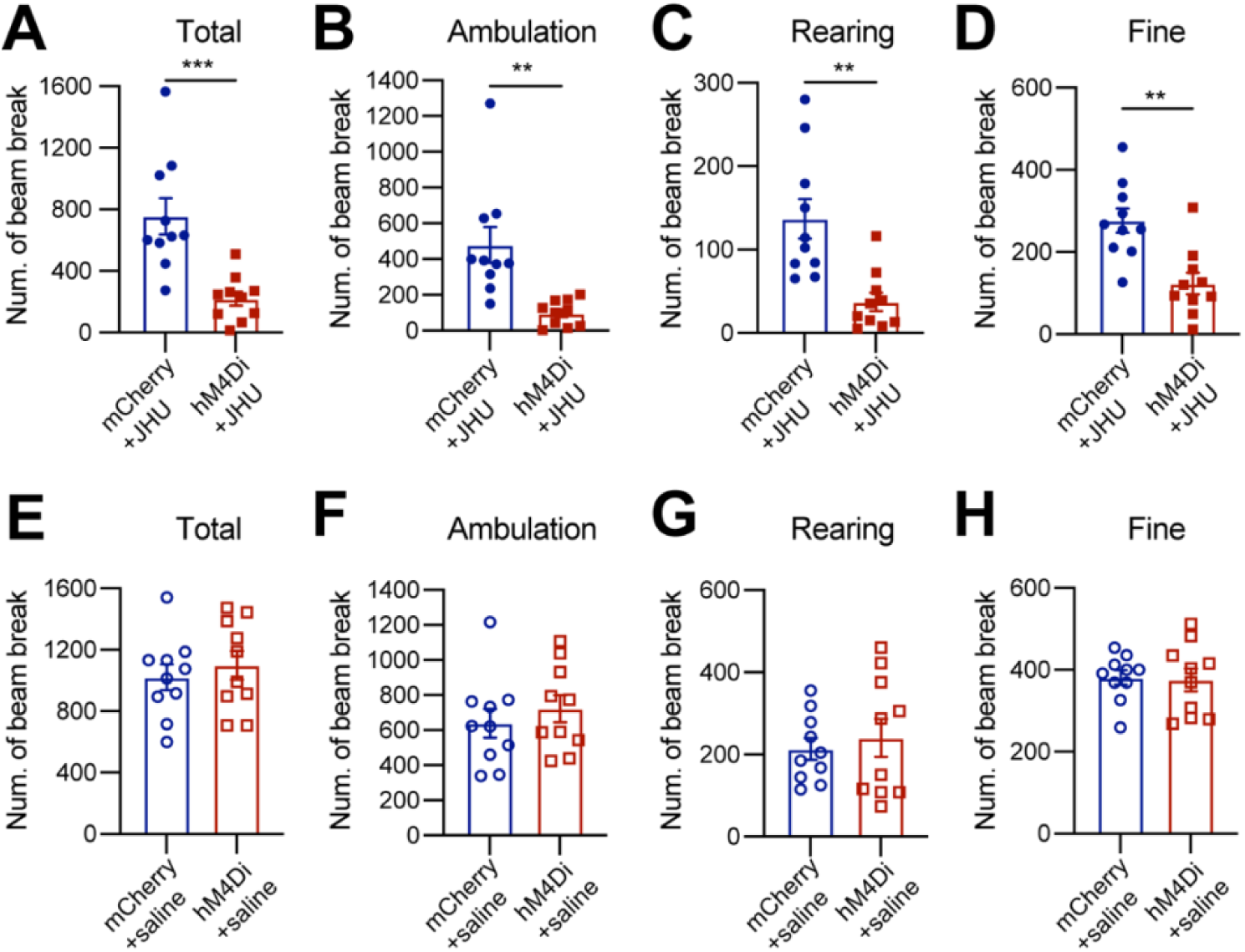
Chemogenetic inhibition of *Calb1*^+^ nigrostriatal DANs impairs spontaneous locomotion. **(A-D)** Quantification of total (**A**) (p = 0.001), ambulatory (**B**) (p = 0.001), rearing (**C**) (p = 0.001), and fine (**D**) (p = 0.001) movement in TC double KI mice injected with either control mCherry (n=10) or hM4Di AAVs (n=10) in the SNc. Locomotor activity was measured during a 30-minute open-field test following i.p. injection of JHU37160. Data are presented as mean ± SEM. unpaired t-test. **(E-H)** Quantification of spontaneous locomotion: total (**E**) (p = 0.540), ambulatory (**F**) (p = 0.435), rearing (**G**) (p = 0.616), and fine (**H**) (p = 0.862) movement with the same cohort of mice as in (**A-D**) following saline injection. Data are presented as mean ± SEM. unpaired t-test.

### Chemogenetic inhibition of *Calb1*^+^ SNc DANs severely impairs motor skill learning

To investigate the role of *Calb1*^+^ SNc DANs in motor skill learning, we conducted rotarod motor skill learning tests ^16,22^ on TC mice that received bilateral injections of control and DREADD AAVs into the SNc. Chemogenetic inhibition with JHU37160 severely impaired motor skill acquisition in hM4Di-expressing mice compared to mCherry-expressing control mice **(Fig. 3)**.

**Fig. 3.**
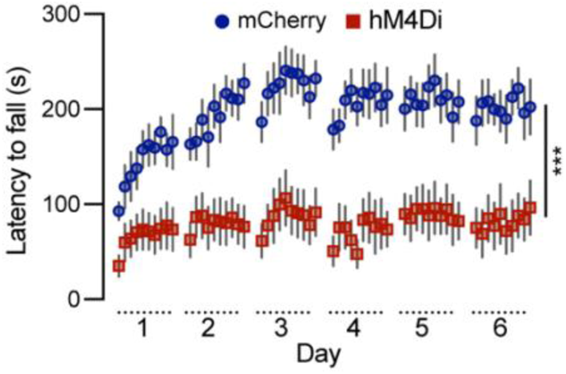
Chemogenetic inhibition of *Calb1*^+^ nigrostriatal DANs severely impairs rotarod motor skill learning. Rotarod motor skill learning in TC double KI mice injected with either control mCherry (n=10) or hM4Di AAVs (n=10) in the SNc. Mice underwent one training session per day for six consecutive days, with JHU37160 administrated prior to each session. Data are presented as mean ± SEM. Two-way ANOVA, mCherry vs. hM4Di effect: *F* (1, 18) = 22.2, *p* = 0.0002.

These data demonstrate that *Calb1*^+^ SNc DANs play a crucial role in acquiring skilled movements.

### Chemogenetic inhibition of *Calb1*^+^ SNc DANs affects the reward learning

DANs are involved in various non-motor function, including reward-related behavior such as seeking rewards, avoidance, and motivation ^10,23,24^. However, the specific role of the *Calb1*^+^ SNc DAN subtype in reward-related behavior remains unclear. To address this, we investigated their involvement in simple reward-related functions using a home cage-based feeding experimentation device (FED3) in food-restricted condition ^25^. We assessed performance in both free-feeding, where a food pellet immediately dispensed following the previous pellet retrieval, and fixed ratio 1 (FR1) reinforcement, where a correct nose-poke to the designated port was required to earn a pellet, schedules. During a 2-hour free-feeding session, DREADD experimental mice and control mice collected a similar number of pellets **(Fig. 4A)**, indicating no difference between the two groups in pellet consumption when pellets were easily accessible.

**Fig. 4.**
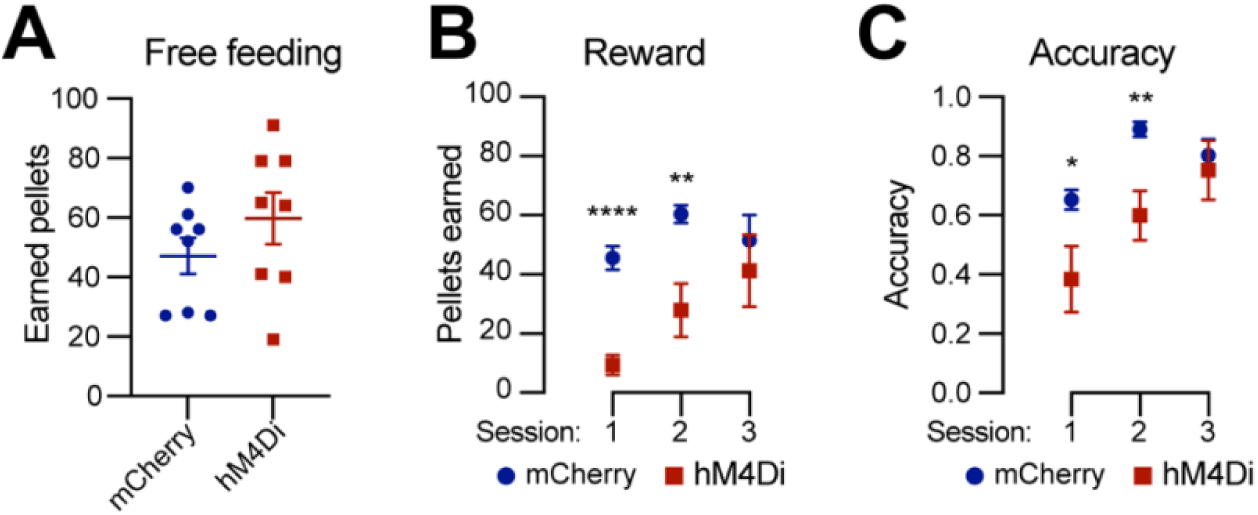
Chemogenetic inhibition of *Calb1*^+^ nigrostriatal DANs affects food-seeking behavior. **(A)** Quantification of rewards earned by control mCherry (n=8) and hM4Di (n=8)-expressing TC double KI mice during a 2-hour free-feeding session on the FED3 device. Data are presented as mean ± SEM. unpaired t-test, p = 0.251. **(B)** Quantification of pellets earned in FR1 training on FED3 across three sessions. Two-way ANOVA with Tukey’s multiple comparison test: p<0.0001 (session 1), 0.0086 (session 2), and 0.5036 (session 3). **(D)** Quantification of nose-poking accuracy in FR1 training on FED3 across three sessions. Two-way ANOVA with Tukey’s multiple comparison test: p=0.0495 (session 1), 0.0096 (session 2), and 0.6679 (session 3).

However, in the FR1 schedule, where mice needed to learn an action-outcome association to obtain a pellet, over first three days, DREADD mice initially exhibited a lower pellet acquisition following chemogenetic inhibition of *Calb1*^+^ SNc DANs but gradually caught up with pellet acquisition rate of control mice **(Fig. 4B)**. Additionally, *Calb1*^+^ DAN-inhibited mice showed reduced accuracy in poking the assigned ports during the first two training sessions but gradually improved and achieved similar accuracy in the subsequent session **(Fig. 4C)**, and remained the same performance level to controls past day 3 (data not shown). These findings suggest that *Calb1*^+^ SNc DANs contribute to early stages of associative learning and reward-seeking behavior, though their influence diminishes over time, leading to comparable performance between experimental and control groups in later stages.

### Selectively targeting of *Aldh1a1*^+^ SNc DANs using intersectional mouse genetics and chemogenetic approaches

Compared to the chemogenetic inhibition of *Calb1*^+^ SNc DANs, genetic ablation of *Aldh1a1*^+^ SNc DANs in a previous study resulted in only a modest impairment of locomotion ^16^. To directly compare the roles of *Aldh1a1*^+^ and *Calb1*^+^ SNc DANs on locomotion within the same experimental paradigm, we generated *Th*^Flp^; *Aldh1a1*^CreERT^^2^ (TA) double KI mice and selectively inhibited *Aldh1a1*^+^ SNc DAN activity using the same chemogenetic approach applied to *Calb1*^+^ SNc DANs **(Fig. 5A)**.

**Fig. 5.**
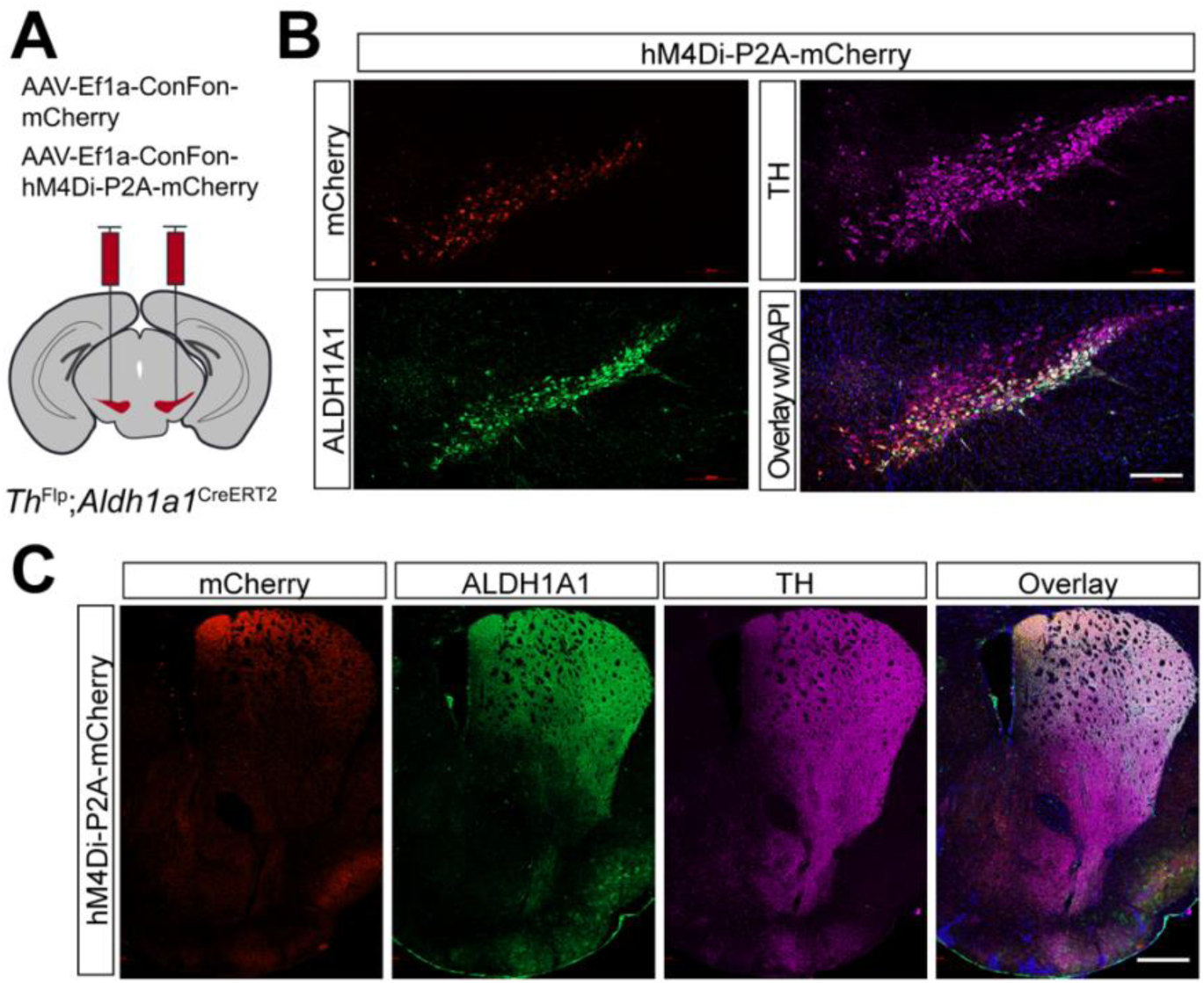
Targeted expression of the chemogenetic inhibitor hM4Di in *Aldh1a1*^+^ nigrostriatal DANs. **(A)** Schematic depicting the stereotactic delivery of control mCherry or hM4Di AAVs in the SNc of *Th*^Flp^; *Aldh1a1*^CreERT2^ (TA) double KI mice. (**B**) Representative images showing mCherry (red), ALDH1A1 (green), and TH (magenta) expression in the SNc of control mCherry (left panel) and hM4Di (right panel)-expressing mice. Scale bar: 1mm. (**C**) Representative images illustrating mCherry (red), ALDH1A1 (green) and TH (magenta) staining in the striatum of control mCherry-expressing mice. Scale bar: 1mm.

Immunostaining confirmed selective expression of hM4Di and mCherry in approximately 44% of *Aldh1a1*^+^ SNc DANs, which are primarily located in the ventral tier of the SNc **(Fig. 5B)**. Examination of their projection patterns in the dorsal striatum revealed axon terminals distributed in both the DMS and dorsolateral striatum (DLS) **(Fig. 5C)**. Together, the use of TA double KI mice combined with stereotaxic injection allows selective targeting of *Aldh1a1*^+^ SNc DANs for genetic and functional manipulations.

### Chemogenetic inhibition of *Aldh1a1*^+^ SNc DANs suppresses spontaneous movements

Using an open-field test to assess spontaneous movements following intraperitoneal injection of the DREADD activator JHU37160, we observed a marked reduction in total movement, ambulatory movement, rearing, and fine movement **(Fig. 6A-D)**. These findings reveal a critical acute role of *Aldh1a1*^+^ SNc DANs in regulating spontaneous movements.

**Fig. 6.**
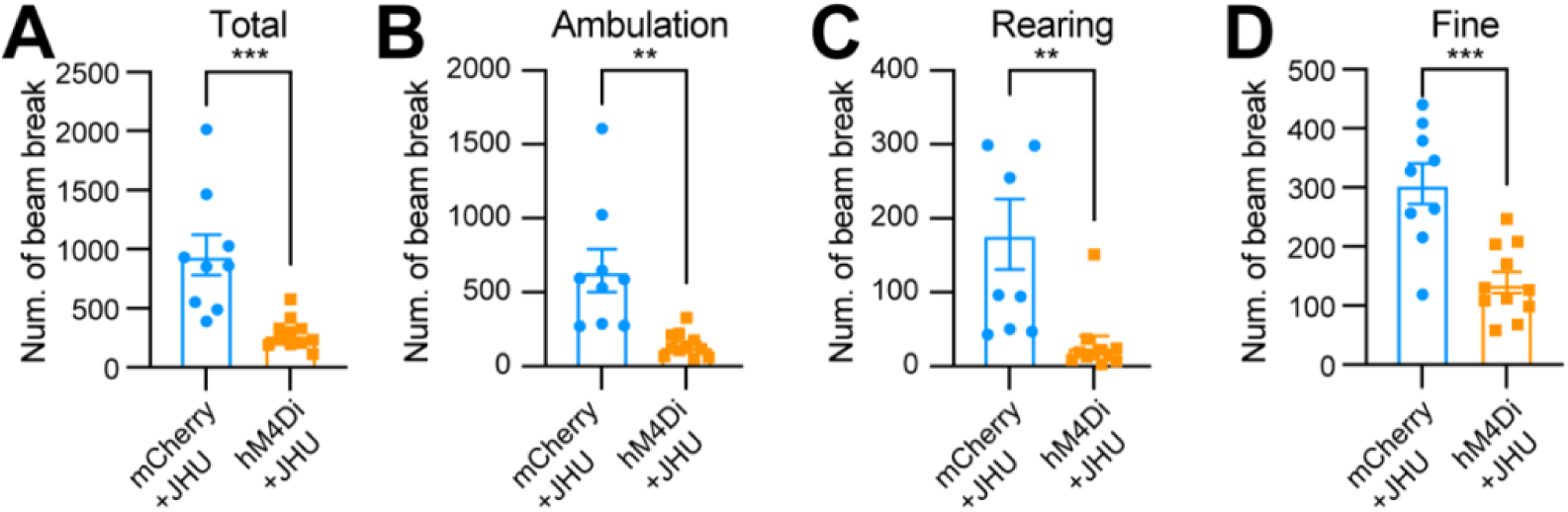
Chemogenetic inhibition of *Aldh1a1*^+^ nigrostriatal DANs impairs spontaneous locomotion. **(A-D)** Quantification of total (**A**) (p = 0.0006), ambulatory (**B**) (p = 0.0013), rearing (**C**) (p = 0.0036), and fine (**D**) (p = 0.0002) movement in TC double KI mice injected with either control mCherry (n=9) or hM4Di AAVs (n=11) in the SNc. Locomotor activity was measured during a 30-minute open-field test following i.p. injection of JHU37160. Data are presented as mean ± SEM. unpaired t-test.

### Chemogenetic inhibition of *Aldh1a1*^+^ SNc DANs severely impairs motor skill learning

To examine the role of *Aldh1a1*^+^ SNc DANs in motor skill learning, we conducted rotarod tests on TA double KI mice that received bilateral injections of control and hM4Di AAVs into the SNc. Chemogenetic inhibition with JHU37160 severely impaired motor skill acquisition in hM4Di-expressing mice compared to mCherry-expressing control mice **(Fig. 7)**. These results confirm previous findings in *Aldh1a1*^+^ SNc DAN-ablated mice ^16^ and further demonstrate that *Aldh1a1*^+^ SNc DANs are essential for acquiring skilled movements.

**Fig. 7.**
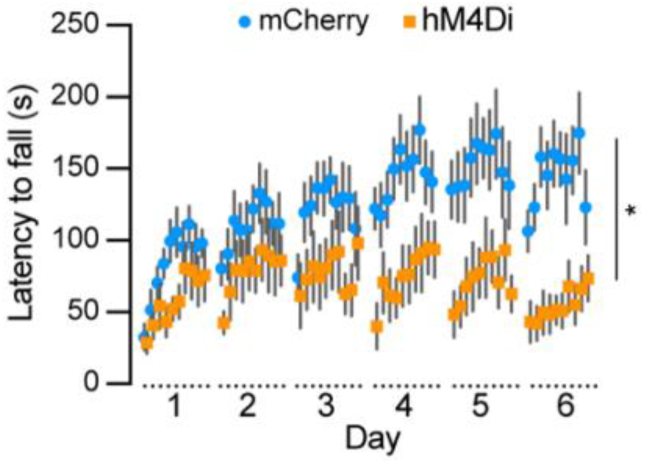
Chemogenetic inhibition of *Aldh1a1*^+^ nigrostriatal DANs severely impairs rotarod motor skill learning. Rotarod motor skill learning in TA double KI mice injected with either control mCherry (n=9) or hM4Di AAVs (n=11) in the SNc. Mice underwent one training session per day for six consecutive days, with JHU37160 administrated prior to each session. Data are presented as mean ± SEM. Two-way ANOVA, mCherry vs. hM4Di effect: *F* (1, 18) = 8.227, *p* = 0.012.

### Chemogenetic inhibition of *Aldh1a1*^+^ SNc DANs does not affect the simple reward learning

To determine whether *Aldh1a1*^+^ SNc DANs are involved in reward and associative learning related behaviors, we examined the performance of *Aldh1a1*^+^ DAN-inhibited mice in both free-feeding and FR1 reinforcement schedules. Chemogenetic inhibition of *Aldh1a1*^+^ SNc DANs had no defect on free feeding, reward acquisition, or nose-poking accuracy **(Fig. 8A-C)**. These findings suggest that *Aldh1a1*^+^ SNc DANs do not play a significant role in the simple associative learning and reward-seeking behaviors, in contrast to that of *Calb1*^+^ SNc DANs.

**Fig. 8.**
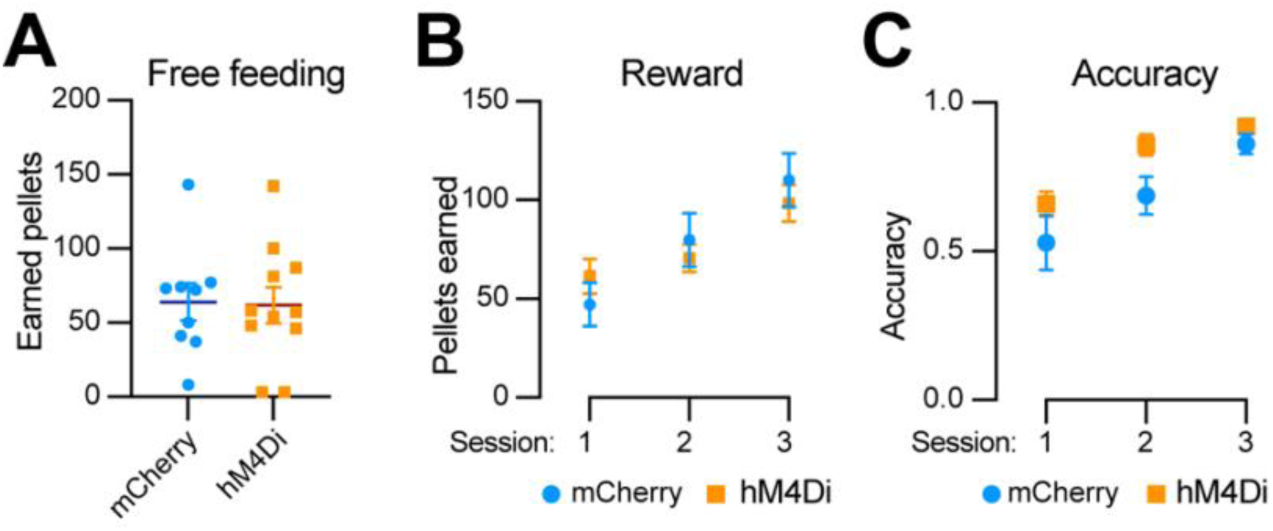
Chemogenetic inhibition of *Aldh1a1*^+^ nigrostriatal DANs does not affect food-seeking behavior. **(A)** Quantification of rewards earned by control mCherry (n=9) and hM4Di (n=11) AAV-injected TA double KI mice during a 2-hour free-feeding session on the FED3 device. Data are presented as mean ± SEM. unpaired t-test, p = 0.251. **(B)** Quantification of pellets earned in FR1 training on FED3 across three sessions. Two-way ANOVA with Tukey’s multiple comparison test: p<0.0001 (session 1), 0.0086 (session 2), and 0.5036 (session 3). **(D)** Quantification of nose-poking accuracy in FR1 training on FED3 across three sessions. Two-way ANOVA with Tukey’s multiple comparison test: p=0.0495 (session 1), 0.0096 (session 2), and 0.6679 (session 3).

## Discussion

In this study, we employed intersectional genetics and a chemogenetic approach to selectively manipulate the activity of *Calb1*^+^ and *Aldh1a1*^+^ nigrostriatal DAN subtypes in TC or TA double KI mice during Open-field, rotarod motor skill learning, and FED3 reward learning tests. We found that inhibition of either DAN subtype suppressed locomotion and impaired motor skill learning, while inhibition of the *Calb1*^+^ subtype also slowed down reward-based reinforcement learning. These findings demonstrate that both DAN subtypes in SNc are essential for promoting locomotion and motor skill learning, with *Calb1*^+^ neurons additionally contributing to reward associative learning.

We previously demonstrated that genetic ablation of *Aldh1a1*^+^ nigrostriatal DANs resulted in a modest reduction in locomotor activity but completely abolished rotarod motor skill learning ^16^. Interestingly, while chemogenetic inhibition of *Aldh1a1*^+^ DANs similarly caused severe motor skill learning deficits, it led to a more pronounced reduction in locomotor activity compared to genetic ablation. The underlying reason for this discrepancy remains unclear; however, we speculate that the rapid and transient inhibition of *Aldh1a1*^+^ DAN activity via chemogenetics may not trigger a robust compensatory response from other neural circuits, unlike genetic ablation, which gradually leads to neuronal loss over weeks ^16^.

While the locomotion-promoting function of *Aldh1a1*^+^ nigrostriatal DANs is well established ^16,17^, the role of the *Calb1*^+^ subtype remains less defined. A recent study reported that *Calb1*^+^ DAN activity correlates with deceleration, as measured by calcium transients in axons or somas of specific DAN subtypes while head-fixed mice ran on a treadmill ^17^. This correlation with deceleration suggests a potential role for *Calb1*^+^ DANs in locomotion suppression. However, using chemogenetic manipulation in freely moving mice, we demonstrated that inhibition of *Calb1*^+^ DAN activity led to a severe reduction in locomotor activity, consistent with the general role of basal ganglia dopamine transmission in promoting movement.

Furthermore, to explore the involvement of *Calb1*^+^ DANs in reward-related behaviors, we employed the FED3 system. Our findings revealed that *Calb1*^+^ DANs contribute to early associative-learning and reward-seeking behaviors, as chemogenetic inhibition disrupted food pellet acquisition and performance accuracy, particularly during the initial phase of the FR1 sessions. These results align with previous studies highlighting the role of *Calb1*^+^ nigrostriatal DANs in reward processing ^17^.

It is worth noting that a small fraction of DANs located in the ventromedial region of the SNc co-express both *Aldh1a1* and *Calb1*. These neurons project their axonal fibers primarily to the DMS. However, their exact roles in motor control, learning, and reward responses remain unknown. Given that both *Aldh1a1*^+^ and *Calb1*^+^ DAN subtypes promote locomotion and motor learning, it is likely that this dual-positive subtype also facilitates these functions. However, future studies employing additional intersectional genetic manipulations will be necessary to specifically investigate the functional contributions of this distinct DAN population.

Advances in single-cell and single-nucleus RNA sequencing have provided valuable insights into the complexity and heterogeneity of DAN subtypes in the SNc ^10,11,26–28^. By leveraging the distinct molecular markers identified in these neuronal subtypes, our study highlights the importance of systematically defining their functions. This approach will enhance our understanding of the diverse motor and non-motor phenotypes observed in PD.

## MATERIALS AND METHODS

### Mouse work

All mouse studies were conducted in accordance with the guidelines approved by the Institutional Animal Care and Use Committees (IACUC) of the Intramural Research Program of the National Institute on Aging (NIA), NIH. *Th*^Flp^ mice (strain: C57BL/6N-Thtm1Awar/Mmmh; Stock# 050618-MU; MMRRC item: 050618-MU-HET) were obtained from the Mutant Mouse Reginal Resource Center (MMRRC). *Calb1*^IRESCre^ mice were obtained from the Jackson Laboratory (Bar Harbor, ME, USA; JAX# 028532). Mice used for viral injections were between 2 and 4 months of age. They were housed in groups of 2-5 animals under a 12-hour light/12-hour dark cycle with *ad libitum* access to water and a standard diet. Both male and female mice were included in the experiments, and all behavioral tasks were conducted during the light cycle.

Littermates were randomly assigned to different experimental groups before the start of the study.

### AAV Vectors

The AAV-EF1a-Con/Fon-mCherry vector ^29^ was obtained from Addgene (#137132, Watertown, MA, USA). The AAV-EF1a-Con/Fon-GiDREADD-P2A-mCherry vector was constructed based on the pAAV-nEF-Con/Fon-ChR2-mCherry plasmid ^29^. The DNA cassette was designed by splitting the reading frame of hM4Di-P2A-mCherry into three fragments, which were synthesized *de novo* and incorporated into the pAAV-EF-Con/Fon-GiDREADD-P2A-mCherry vector.

### Animal surgery and stereotaxic injections

We selected adult male and female mice, aged 20-24 weeks and weighing >20 grams, for stereotactic injection as described previously ^30^. In brief, mice were deeply anesthetized with isoflurane, and the head fur was shaved and cleaned with iodine solution. The anesthetized mice were then placed on a stereotactic frame (Stoelting) equipped with stabilizing ear bars and a mouthpiece delivering a steady flow of 1.5% isoflurane and 2% oxygen. The levels of oxygen and isoflurane were continuously monitored throughout the procedure. A midline scalp incision was made to expose the skull, and Bregma and Lambda coordinates were used for precise brain positioning. A small hole was drilled over the target region using a 0.8 mm stereotaxic drill.

Injection coordinates for the midbrain were determined based on the standard mouse brain atlas. Following the injection, the needle remained in place for five minutes to allow viral diffusion and prevent backflow along the needle tract upon withdrawal. The scalp was then closed using non-absorbable sutures and Vetbond tissue adhesive (3M), and an antibiotic ointment was applied. 350 nL of AAV vectors (original titer: ∼9 × 10^13 VG/mL, diluted 1:3 in sterile PBS) were bilaterally injected into the SNc at the following coordinates from Bregma: AP –3.10 mm, ML ±1.2 mm, DV –3.8 mm. The injections were performed using a syringe controlled by a motorized stereotaxic injector at a rate of 75 nL/min.

### Brain sectioning and Immunohistochemistry

Mice were anesthetized with pentobarbital and transcardial perfusion was performed with a controlled pressure device with 100-150 ml ice cold 1x PBS followed by 100 ml 4% paraformaldehyde (PFA) in 1x PBS. Brains were carefully removed and post-fixed in 4% PFA in 1x PBS overnight at 4^0^C. Brains were then washed in PBS for 1 min and transferred to cryo-protectant 30% sucrose in 1x PBS solution for 2 days at 4^0^C (until the brain sank). Dehydrated Brain tissues were cut into 35-40 µm thick coronal sections using Leica cryostat (Leica Biosystems) unless specified otherwise. Brain sections were collected in a 24 well plate in 1x PBS with 0.01% sodium azide (sigma Aldrich) and stored at 4^0^C.

For immunohistochemistry, 35-40 µm thick free-floating coronal sections were washed in one time for 10 min in 1x PBS briefly, followed by a for1 h at room temperature in blocking solution containing 10% bovine serum albumin (BSA) and 0.3% Triton X-100 in PBS. Primary antibodies were freshly prepared in diluted career solution followed by overnight incubation with gentle shaking at 4°C.‘Brain sections were then washed with 3 x10 min with career solution. The sections were then incubated with Alexa Fluor-conjugated secondary antibodies diluted in (1:500) in career solution against the specific host of primary antibodies at room temperature for 1 hr. The brain sections were then washed in carrier solution for 2 X 10 minutes. for Nuclei were stained with DAPI (Sigma, 1:1000) in PBS for 1 min followed by washed in 1x PBS for 10 min. The stained sections were mounted onto slides and cover slipped with hard setting ProLong Gold Antifade liquid mounting media (Invitrogen # P36930). The following antibodies were in our experiments, anti-ALDH1A1 (1:500, polyclonal goat IgG, R&D systems # A5869), anti-tyrosine hydroxylase (1:1000, polyclonal rabbit IgG, Sigma-Aldrich # AB152), anti-RFP (1:3000, chicken IgG, Aves Lab).

### Confocal microscopy and image analysis

Fluorescent staining images were acquired using a Zeiss LSM 780 and LSM 980 confocal microscopes (Zeiss). Whole coronal sections were initially captures using a 10× objective lens. Higher magnification images of regions of interest were then obtained with a 20× objective lens using z-stack intervals. The 10× images were 512 × 512 pixels, while all 20× images were at least 1024 × 1024 pixels. Counts were taken from comparable sections across the rostro-caudal axis of the midbrain. Imaging was performed using three to four fluorescence channels with the following filters: Alexa Fluor 405, 488, 555, and 647. Each channel was imaged separately to eliminate potential signal crosstalk between channels.

### Cell counting

Two coronal brain sections containing the *substantia nigra pars compacta* (SNc) were selected from each mouse (n = 5 *Aldh1a1*^CreERT2^; *TH*^Flp^ and 3 *Calb1*^Cre^; *TH*^Flp^). RFP, ALDH1A1, and TH cells were quantified per section using NeuroInfo analysis software (MBF Bioscience).

Cell detection was performed using the “Detect Cells” pipeline, and regions of interest (ROIs) were manually delineated around the SNc using TH immunostaining and the Allen Brain Atlas as anatomical references. ROIs were drawn by placing connected contour points with the mouse and closed by selecting “Close Contour”. The previously drawn ROI was then selected to proceed to the preview and detection step. Channel-specific detection parameters were defined as follows: large diameter = 25.00 µm, small diameter = 8.00 µm, and detection strength = 1000.

Following automated detection, each section was reviewed and manually corrected for false-positive or false-negative labeling. This detection procedure was performed independently for each of the three channels. Colocalization analysis was conducted by right-clicking on the marker icon of a given channel and selecting “Colocalize Markers”. Marker pairs and a new marker for colocalization were defined, the “Delete marker combination when placing a colocalization marker” option was deselected, and the colocalization distance threshold was set to 8 µm. Cell counts were exported to Excel, and average values were calculated for each section. Group averages were then used to assess viral transduction efficiency across cohorts.

### DREADD ligand JHU37160 Drugs preparation

JHU37160-dihydrochloride (Cat# HB6261, Hello Bio, Inc. Princeton, NJ, USA), a high-affinity and highly potent water-soluble DREADD agonist ^21^, is stored as powder at −20°C. A new 5 mg bottle was dissolved in 16.666 mL of sterile 0.9% sodium chloride solution to achieve a final concentration of 0.3 mg/mL. The solution was aliquoted into 5 mL Eppendorf tubes, labeled, and stored at −20°C for up to one month. On the day of injection, one tube was thawed and diluted 10-fold with sterile saline before use. Mice received intraperitoneal injections of JHU37160 (0.3 mg/kg bodyweight) 30 minutes before each motor or feeding test.

### Behavioral assessment Habituation of animals

For all behavioral studies, mice were housed in groups, with males and females housed in separate cages. Mouse cages were transported on a cart covered with lab coat from the animal home room to the rodent behavior core. Mice were then habituated in a small holding room for 20-30 minutes before starting the behavior. The paradigm was cleaned with 70% alcohol and dried before placing the animal in the designated space.

### Motor coordination and activity tests Open-field test for spontaneous locomotion

To measure the motor activity, we used the Photobeam Activity System based Open field (PAS-OF, San Diego Instruments) test. The chambers were cleaned with 70% alcohol, turned on red lights, and placed mice in their designated chambers according to their IDs. The mice explored the chambers, and their movements were recorded by two sets of 4 × 8 photobeams for 30 minutes. Infrared photo beams at two different heights tracked the mice movements: the lower beams recorded x- and y-axis positions, termed fine and ambulation movements, while the upper beams counted rearing or vertical movements. The PAS-OF system accurately recorded all beam interruptions caused by the mice movements including ambulation, fine, and rearing behavior. It utilizes a database to store all results in a single Microsoft Access database file in table format easy to export as CSV format. We then run the analysis program R scripts to extract the different parameters of locomotion. The system recorded any activity as ambulatory movement when three or more different light beams were interrupted consecutively, whereas it was recorded as fine movement when only adjacent light beams were disrupted alternately.

### Rotarod test for motor skill learning

One week after the open field activity test, we assessed the motor skill learning of mice using accelerating rotarod tests, following the similar procedure we’ve done in the past ^16^. We cleaned the 5-lane rotarod instrument (Pan lab, Harvard Apparatus, Holliston, MA) with 70% alcohol and kept the rod dry. We then gently placed the mice on the rotating rod and gradually increased the speed linearly from 4 to 40 revolutions per minute over a 5-minute trial. The duration the mice stayed on the rod was recorded as latency to fall. Ten trials were conducted per day for six consecutive days, with a 2–3-minute interval between each trial.

### Operant conditioning using FED3

#### Feeding experimentation device 3

The Feeding Experimentation Device 3 (FED3) is a compact battery-powered operant device designed for feeding and training rodents within their home cage environment ^25^, purchased fully-assembled from Open Ephys (Open Ephys Production Site, Lisbon, Portugal). It features a dual nose-poke system with left and right sensors, a beeper, a multi-color LED strip, a pellet dispenser that dispenses pellet, an output jack for equipment connection, a pre-loaded ClassicFED3 program, and a user communication screen. All behavioral activity interacting with the device is logged onto an onboard microSD card for future analysis.

#### Food restriction

Rodents were food-restricted overnight one to two days before starting the feeding device experiment. During the restriction period, body weight, eating time, and reward intake were recorded daily and maintained in a log accessible to the facility veterinarian in the animal holding room. If an animal’s body weight dropped below 85% of its initial weight, it was given unrestricted access to food and monitored until full recovery. Mice were individually housed into a new cage containing a FED3 device for a 2-hour training session period. The pellets dispensed from the FED3 device were the only food source available during the training session. After training, they were returned to their original group home cage and given unlimited access to regular chow for three hours, starting 30 minutes after the session ended.

#### Free feeding

A FED3 device, half-filled with grain-based dustless pellet reward (Bio-Serv, 20 mg, Product # F0163, Flemington, NJ, USA), was placed in a rodent’s cage. The device was powered on and set to “free feeding” mode, allowing mice to retrieve pellets in the food magazine without requiring a nose poke. Half hour following either JHU37160 or saline injections, mice were placed in the new cage and given access to the FED3 device for a 2-hour session. In this free-feeding paradigm, pellet availability was independent of nose-poking behavior. The pellet sensor detected when a pellet was removed from the magazine and automatically dispensed another pellet. After one or two sessions, mice became habituated to the FED3 pellet-dispensing system and were transitioned to fixed-ratio training.

#### Fixed ratio 1 (FR1) schedules of reinforcement

Mice transitioned from free-feeding mode to a fixed-ratio 1 (FR1) reinforcement training schedule, where they received pellets through active nose-poking in a similar experimental setting with FED3 device and single housing. Each experiment started half hour following either JHU37160 or saline injections. The device was powered on and set to “FR1” mode. During FR1 sessions, a nose poke in the “active” port triggered the immediate delivery of a single 20mg pellet reward in the magazine, while a nose poke in the “inactive” port did not result in pellet delivery. In all experiments, the left nose poke was designated as the active port, while the right nose poke was designated as the inactive port. Correct nose-poking behavior was reinforced with immediate audio and visual cues. There was no punishment after poking inactive port. To assess the reinforcement learning behavior, mice were trained on consecutive days with 2-hour FR1 sessions. Performance data from the first to the third sessions were used for the analysis.

Accuracy was calculated as the count of pokes on active port divided by total poke count in a session. Mice were trained with FR1 task for 3-7 sessions.

### Statistics

Data were analyzed using Prism 10 (GraphPad) and custom R code. Statistical details for each experiment are provided in the figure legends. Results are presented as means ± SEM. Statistical significance was assessed using unpaired parametric t-tests and two-way ANOVA with multiple comparisons. Significance defined as *p < 0.05, **p < 0.01, ***p < 0.001, and ****p < 0.0001.

## Acknowledgements

This work is supported in part by the Intramural Research Programs of National Institute on Aging, NIH (HC, ZIA AG000944, AG000928), the Rodent Behavioral Core of Intramural Research Program of National Institute of Mental Health (MH002952), and National Nature Science Foundation of China (WDL, 32220103006). We thank members of Cai lab and the Rodent Behavioral Core of Intramural Research Program of National Institute of Mental Health for their suggestions and technical assistance.

## Author contributions

H.C. conceived and wrote the manuscript and prepared the figures with inputs from all authors. A.H. designed and performed histology, behavioral experiments and data analyses, prepared figures, wrote methods and figure legends, and wrote the manuscript for the TC double KI mice. G.R. designed and conducted performed histology, behavioral experiments and data analyses for the TA double KI mice. L.T. performed histology. D.B. generated the cohort of TA mice. L.S. built the dual control hM4Di plasmid. L.C. performed stereotactic surgery. V.M. contributed to histological analysis. L.W. designed FED3 experiments, contributed to data analysis, and edited the manuscript. W.L. edited and provided critical feedback. All authors read and approved the final manuscript.

